# p75NTR modulation by LM11A-31 counteracts oxidative stress and cholesterol dysmetabolism in a rotenone-induced cell model of Parkinson’s disease

**DOI:** 10.1101/2024.12.25.630306

**Authors:** Daniele Pensabene, Noemi Martella, Giuseppe Scavo, Michela Varone, Emanuele Bisesto, Francesca Cavicchia, Mayra Colardo, Sandra Moreno, Marco Segatto

**Author notes:** Correspondence (M.S.), (D.P.).

## Abstract

Neurotrophins play pivotal roles in the development and proper functioning of the nervous system. The effects of these growth factors are mediated through the binding of high-affinity receptors, known as Trks, as well as the low-affinity receptor p75NTR. The latter has the capacity to induce complex signal pathways, as it favors both pro-survival and pro-apoptotic cascades depending on the physiopathological condition. Recent findings have indicated that p75NTR expression is increased in post-mortem Parkinson’s disease (PD) brains, and this upregulation is associated with a significant reduction in neuroprotection. Given its double-edged sword nature, p75NTR has recently been identified as a promising therapeutic target to counteract neurodegenerative events. The present study aims to assess the neuroprotective effects of p75NTR modulation in a rotenone-induced neuronal model of PD. To this end, differentiated SH-SY5Y cells were exposed to rotenone to mimic the PD phenotype, and the small molecule LM11A-31 was used to modulate p75NTR activity. The main results revealed that LM11A-31 significantly mitigated the hallmarks of PD, including cell death, neuromorphological aberrations, and α-synuclein accumulation. Pharmacological manipulation of p75NTR also reduced oxidative damage by increasing the expression of transcriptional factors that regulate the antioxidant response and by decreasing the expression of the pro-oxidant NADPH-oxidase modulatory subunits. Furthermore, LM11A-31 hampered cholesterol buildup induced by rotenone, normalizing the expression of proteins involved in cholesterol biosynthesis, uptake and intracellular trafficking. Taken together, these findings suggest that p75NTR modulation may represent a novel approach to counteracting PD abnormalities of redox and cholesterol metabolism.

## Introduction

p75NTR is the first member discovered of the tumor necrosis factor (TNF) receptor family (Colardo et al., 2021). The activation of this receptor is mediated by the unselective binding to all neurotrophins (NTs), a small group of growth factors involved in the regulation of both development and function of the nervous system, modulating the balance between neuronal survival and cell death (Bucci et al. 2014; Pallottini et al. 2020). Both precursor and mature neurotrophins exert biological functions by binding p75NTR. Specifically, mature neurotrophins bind p75NTR with low affinity, determining cellular survival and neurite outgrowth; whereas NTs precursors preferentially bind p75NTR favoring apoptosis under physiological conditions (Segatto et al. 2019; Longo et al. 2013). For this reason, p75NTR has emerged as a pivotal mediator of both survival and cell death, depending on the interaction with various ligands and co-receptors: for instance, alterations in its expression or atypical processing may result in extensive cell death, thus contributing to the onset and development of neurodegenerative diseases such as Alzheimer’s disease (AD) (Zeng et al. 2011), Huntington’s disease (HD) (Simmons et al. 2017) and Amyotrophic Lateral Sclerosis (ALS) (Shu et al. 2015). Additionally, several lines of evidence demonstrated that p75NTR signaling regulates dopaminergic cell survival and death under physiological conditions, suggesting a crucial involvement of this receptor in the pathogenesis of Parkinson’s disease (PD) (Chen et al. 2008; Ali et al. 2023).

PD is the first most common neurodegenerative disease affecting movement, with patients displaying bradykinesia, resting tremors and overall loss of motor control. These alterations depend on the extensive dopaminergic cell death in the *Substantia Nigra pars compacta* (*SNpc*), a region of the brain necessary for locomotor activity regulation (Liu et al. 2021; Obeso et al. 2017). Despite years of scientific research, PD etiology remains quite elusive. However, various neurodegenerative hallmarks were detected such as cytoplasmatic inclusions of α-Synuclein (α-syn), mitochondrial impairments, oxidative stress, neuroinflammation and autophagy derangements (Ibarra-Gutiérrez et al. 2023; Subramaniam et al. 2013). Additionally, several findings identified cholesterol metabolic disbalances as a less-known pathological hallmark in PD. In fact, autoptic brains from PD patients display alterations of lipid rafts organization on the cell membrane contributing to its disruption (Fabelo et al. 2011). Notably, recent evidence indicated a downregulation of Niemann-Pick C1 (NPC1) protein and a subsequent build-up of unesterified cholesterol within lysosomes in an *in vitro* model of PD. These results suggest the occurrence of a Niemann-Pick type C disease (NPCD)-like phenotype, further highlighting the relevance of cholesterol dysmetabolism in PD (Caria et al. 2023). Unfortunately, a resolutive therapy for PD is not available yet. Currently, dopamine precursor levodopa (L-DOPA) is employed as a therapeutic strategy to increase dopamine concentration in the *striatum*. Nevertheless, L-DOPA treatment is associated with both tolerance over time and drug-related adverse events (Zheng et al. 2021). It is therefore imperative to seek new strategies to rescue the dopaminergic cell death in the *SNpc*. In this context, p75NTR may represent a valuable candidate, as its activation was found to selectively favor dopaminergic neuronal death and downregulate transcription factors entangled with neuroprotection, thereby contributing to the progression of the neurodegeneration (Chen et al. 2008; Alavian et al. 2009; Ali et al. 2023). According to this notion, it has been recently shown that p75NTR overexpression is associated with the severity of dopaminergic cell death in a rat model of PD (Liu et al. 2021).

The double-edged sword nature of p75NTR in driving both cell death and cell survival led to the identification of small molecules capable of modulating the activity of p75NTR to attenuate the severity of neurological disorders. These p75NTR-binding molecules, including THX-B, LM11A-24, and LM11A-31, efficiently suppress the activation of transduction pathways involved in cell death while concurrently enhancing trophic signaling. Particularly, LM11A-31 is specifically designed to bind to p75NTR within the NGF loop1 binding domain and is the best characterized modulator to date. It is neither a strict agonist nor antagonist, rather it allows for pro-survival signaling induced by phosphoinositide 3-kinase (PI3K)/protein kinase B (PKB) while concurrently suppressing degenerative pathways (Massa et al., 2006).

The pharmacological modulation of p75NTR by LM11A-31 has reported promising results in AD and HD animal models and clinical trials, favoring neuronal resilience and slowing the progression of pathophysiological features (Simmons et al. 2017, Nguyen et al. 2022; Shanks et al., 2024). Despite this evidence, there is a paucity of knowledge concerning the pharmacological modulation of p75NTR in PD.

Hence, the aim of this study was to evaluate the beneficial effects of LM11A-31 in an *in vitro* model of PD by exposing SH-SY5Y differentiated cell line to rotenone, a well-known environmental toxin that mimics the pathological phenotype. Particularly, we choose the PD model induced by rotenone exposure as it faithfully reproduces key mechanisms of PD pathogenesis, such as build-up and aggregation of α-syn, progressive oxidative damage, and apoptotic cell death (Sherer et al., 2002).

## 2. Materials and Methods

### 2.1 Cell Cultures

SH-SY5Y human neuroblastoma cell line was cultured at 5% CO2 in DMEM medium at high glucose (D6429, Merck Life Science, Milan, Italy), containing 10% (*v/v*) fetal bovine serum (FBS, F7524, Merck Life Science, Milan, Italy) and 1% penicillin/streptomycin solution (P06-07100, PAN Biotech, Aidenbach, Germany). Throughout the experiments, 250000 cells were seeded; after 5 hours, SHSY5Y differentiation was induced by incubating cells with DMEM, 1% FBS and 10 μM of retinoic acid (R2625, Merk Life Science, Milan, Italy) for 72 hours. Then, cells were pre-treated either with p75NTR modulator LM11A-31 (0,5 μM) or with vehicle (DMSO, 1:1000 dilution in DMEM, FBS 1%). After 24 hours, cells were exposed to rotenone (0,1 μM R8875, Merck Life Science, Milan, Italy) for other 24 hours.

### 2.2 Morphological evaluation

SH-SY5Y cells were cultured in six-well plates and treated as abovementioned. Cells were observed and photographed employing Nikon Eclipse 7S10 optical microscope at a 20x magnification. Neurite length (expressed as arbitrary units), neurite network (sum of neurite lengths in each field) and neurite-bearing cells (expressed as ratio between the number of neurite-bearing cells and total cell number in the field) were analyzed with ImageJ software v.154d for Windows 10 (National Institutes of Health, Bethesda, MD, USA). Afterwards, cells were exposed to trypsin for 3 minutes and detached from the well surface. Samples were collected and cell count was conducted in Fast-Read 102® (BVS100, Bio Sigma) chambers employing Nikon Eclipse 7S10 microscope.

### 2.3 Western blot analysis

Protein lysate and Western blot were performed according with previously published protocols, with modifications (Trapani et al., 2011; Tonini et al., 2020; Tonini et al., 2021). Briefly, SH-SY5Y were sonicated in sample buffer (Hepes 10 mM, KCl 10mM, MgCl2 1.5 mM, NP-40 0,1%, DTT 0,5mM, protease and phosphatase inhibitor cocktail) for 30s to obtain a total cell lysate. Protein concentration was assessed with Lowry’s method and the Laemmli buffer was added for protein denaturation. The samples were boiled at 95 °C for 3 minutes. Twenty micrograms of protein extract were run employing SDS-PAGE electrophoresis. The Trans-Blot Turbo system (Biorad Laboratories, Milan, Italy) was used for protein transfer on nitrocellulose membranes. The membranes were later incubated at room temperature for 1 hour with 5% fat free milk in DPBS (D1408, Merck Life Science, Milan, Italy, pH 7.4) and subsequently incubated with the primary antibody overnight at 4 °C (Table 1). After incubation, membranes were washed with DPBS to discard excessive primary antibody and exposed for 1 hour to HRP-conjugated secondary antibodies. Clarity ECL Western blotting (1705061, Bio-Rad Laboratories, Milan, Italy) was used for protein-antibody complex observation. Images were captured and densitometric analysis was carried out with ImageJ software v.154d for Windows 10 (National Institutes of Health, Bethesda, MD, USA). All samples were normalized for protein loading with GAPDH as housekeeping proteins. Densitometric calculations were expressed in arbitrary units, determined by the ratio between the protein band intensity and the respective housekeeping protein and normalized to the control levels.

**Table 1.** List of antibodies employed in this work.

| Antibody | Ref | Provider | Application |
| --- | --- | --- | --- |
| **4-HNE** | MA527570 | ThermoFisher Scientific | IF 1:100 |
| **8-OH(d)G** | sc-66036 | SantaCruz<br>Biotechnology | IF 1:100 |
| **GAPDH** | sc-32233 | SantaCruz<br>Biotechnology | WB 1:5000 |
| **HMGCR** | NBP2-61616 | Novus | IF 1:300 |
| **LRP1** | sc-25469 | SantaCruz<br>Biotechnology | IF 1:100 |
| **NOX2** | sc-130543 | SantaCruz<br>Biotechnology | WB 1:1000 |
| **NOX4** | sc-518092 | SantaCruz<br>Biotechnology | IF 1:50 |
| **NPC1** | NB400-14855 | Novus | WB 1:2000 |
| **NRF2** | sc-365949 | SantaCruz<br>Biotechnology | IF 1:100 |
| **N-Tyrosine** | sc-32757 | SantaCruz<br>Biotechnology | IF 1:100 |
| **p22<sup>phox</sup>** | sc-130551 | SantaCruz<br>Biotechnology | IF 1:50 |
| **p47<sup>phox</sup>** | sc-17844 | SantaCruz<br>Biotechnology | IF 1:50 |
| **p75NTR** | Sc-271708 | SantaCruz<br>Biotechnology | IF 1:100 |
| **PGC1α** | Sc-13067 | SantaCruz<br>Biotechnology | IF 1:50 |
| **PPARα** | Ab8934 | Abcam | IF 1:300 |
| **PPARγ** | sc-7273 | SantaCruz<br>Biotechnology | IF 1:100 |
| **SQE** | sc-271661 | SantaCruz<br>Biotechnology | WB 1:500 |
| **SQS** | sc-271602 | SantaCruz<br>Biotechnology | WB 1:500 |
| **SREBP2** | sc-13552 | SantaCruz<br>Biotechnology | IF 1:40 |
| **α-synuclein** | sc-12767 | SantaCruz<br>Biotechnology | IF 1:50 |

### 2.4 Immunofluorescence

Immunofluorescence visualization was performed as previously reported with some modifications (Colardo et al., 2023; Martella et al., 2023). Cells were seeded on sterilized and poly-L-lysine (P6282, Merck Life Science, Milan, Italy)-coated coverslips. After treatment, cells were fixed at room temperature either with 4% paraformaldehyde (D1408, Merck Life Science, Milan, Italy) solution in DPBS or with methanol (32215, Merck Life Science, Milan, Italy) for 5-10 minutes. Subsequently, cells were permeabilized for 5 minutes with DPBS containing Triton 0,1% (X100, Merck Life Science, Milan, Italy). To prevent non-specific binding of the antibody, cells were later exposed to a 3% bovine serum albumin (BSA, A3912, Merck Life Science, Milan, Italy) solution in DPBS containing 0,1% Triton at room temperature for 1 hour. Afterwards, cells were exposed overnight to primary antibodies (Table 1). The next day, samples were incubated for 1 hour with either anti-mouse or anti-rabbit fluorescent conjugate antibodies (Table 1). DAPI (D9542, Merck Life Science, Milan, Italy) was used for nuclei staining. Coverslips were later mounted with Fluoroshield mounting medium (F6182, Merck Life Science, Milan, Italy) and visualized through confocal microscopy (TCS SP8, Leica, Wetzlar, Germany) at 40× magnification. Images were acquired with LAS X software (version 3.5.5) (Leica Camera, Wetzlar, Germany) for Windows 10. All acquisition parameters were unchanged across the experimental group. Signal quantification was performed with ImageJ software v.154d (National Institutes of Health, Bethesda, MD, USA) for Windows 10 as ratio of mean fluorescence intensity to cell area, later normalized to the control level.

To evaluate mitochondrial morphology, MitoTracker™ Dyes for Mitochondria Labeling (M7512, Merck Life Science, Milan, Italy) was used according to producer instructions.

### 2.5 Statistical analysis

Results are shown as mean ± standard deviation (SD). Unpaired Student’s t-test was used to compare mean values between two experimental groups. To evaluate three experimental groups, one-way analysis of variance (ANOVA) was performed along with Tukey’s *post hoc* test. p < 0.05 was considered as statistically significant. Statistical analysis was conducted with GraphPad Prism 8.4.2 (GraphPad, La Jolla, CA, USA) for Windows 10.

## 3. Results

### 3.1 LM11A-31 mitigates PD hallmarks in rotenone-treated SH-SY5Y cells

Compelling evidence reveals that p75NTR expression is abnormal in various neurological disorders, including Parkinson’s disease (Ali et al. 2023; Zeng et al. 2011; Simmons et al. 2017). Starting from these premises, we assessed whether p75NTR levels were also altered in the PD cellular model employed in this study. Immunofluorescence analysis showed that 24 hours of rotenone exposure significantly increased p75NTR expression (Figure 1A), supporting its involvement in the *Parkinsonian* neurodegeneration (Ali et al. 2023).

**Figure 1.**
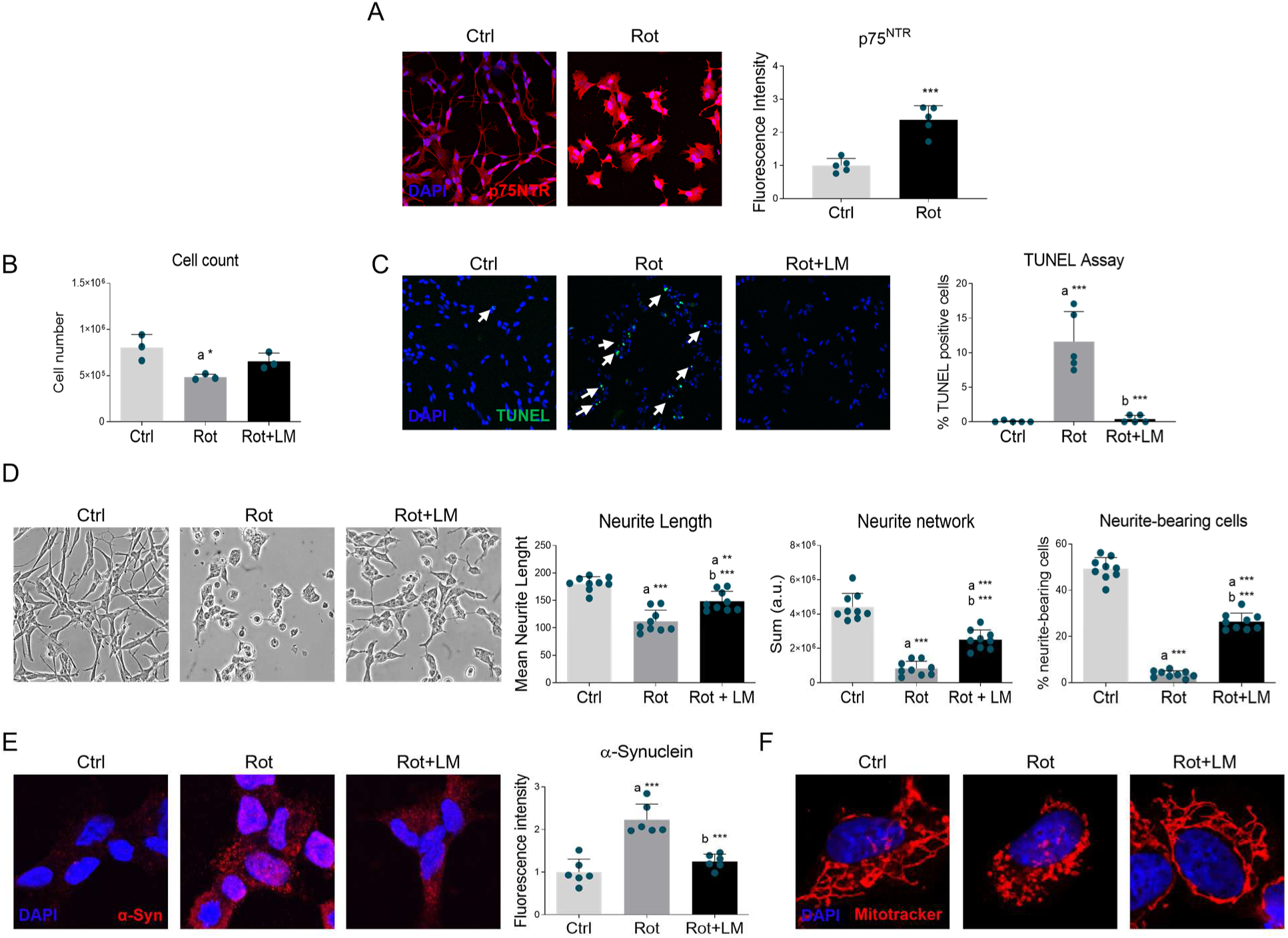
Effects of LM11A-31 on PD hallmarks. (A) Confocal microscopy representative images (left panel) and quantitative analysis (right panel) of p75NTR (red) on differentiated SH-SY5Y cells treated with vehicle (Ctrl) or 100 nM rotenone (Rot) for 24 h. DAPI (blue) was used to counterstain nuclei. N=5. Data are represented as means ± SD. Statistical analysis was performed by using the Unpaired Student’s *t-*test. *** p < 0.001. (B) Cell count in SH-SY5Y differentiated cells treated with vehicle (Ctrl), 100 nM rotenone (Rot) and rotenone with 500 nM LM11A-31 (Rot+LM) for 24 h. N=3. (C) TUNEL assay (green) of SH-SY5Y cells line treated as described above. DAPI (blue) was used to counterstain nuclei; N=5. (D) Representative brightfield images (left panel) and morphological analysis of the neurite length, neurite network and % of neurite-bearing cells (right panel) were conducted using ImageJ software (National Institutes of Health, Bethesda, MD, USA; Sun-Java). N=3. (E) Confocal microscopy representative images of α-synuclein (red) and (F) Mitotracker (red) on SH-SY5Y differentiated cell line treated as previously reported. DAPI (blue) was used to counterstain nuclei. N=5-10. Data are represented as means ± SD. The blue dots around the SD represent the different biological measurements. Statistical analysis was performed by using one-way ANOVA followed by Tukey’s post hoc test. “a” indicates statistical significance vs Ctrl; “b” indicates statistical significance vs Rot group. * p < 0.05, ** p < 0.01, *** p < 0.001.

To further investigate the consequences of p75NTR alterations in PD we used the synthetic modulator LM11A-31. Exaggerated p75NTR expression induced by rotenone was accompanied by a significant reduction in cell number that was attenuated by LM11A-31 administration (Figure 1B). The effect mediated by rotenone on cell number was dependent on apoptosis, as confirmed by the elevation of TUNEL-positive cells. Notably, p75NTR modulation by LM11A-31 completely abolished apoptotic events induced by the neurotoxin (Figure 1C). The effects were not only restricted to cell viability, but also extended to morphological features. Specifically, a dramatic neurite degeneration was observed upon rotenone stimulation, whereas p75NTR modulation partially prevented rotenone-mediated changes in neurite length, neurite network as well as percentage of neurite-bearing cells (Figure 1D). α-Syn overexpression and aggregation are other key pathological hallmarks in PD, being the most abundant components of Lewy’s bodies (Antony et al. 2013). Immunofluorescence analysis revealed that rotenone treatment elicited the build-up of α-Syn *pucta*-like staining, reminiscent of protein aggregation. Conversely, LM11A-31 markedly reduced protein immunoreactivity (Figure 1E).

Mitochondrial dysfunction in dopaminergic neurons is well-described in PD. Specifically, this alteration is well mimicked by rotenone treatment, acting as a potent and reversible mitochondrial complex I inhibitor (Ibarra-Gutiérrez et al., 2023). Mitotracker analysis did not reveal any significant variation in mitochondrial mass. However, according to previous reports (Fang et al., 2013; Ahmad et al., 2013), mitochondrial network was found to be markedly fragmented in rotenone-treated SH-SY5Y. Specifically, control cells displayed branched and tubular mitochondria, whereas rotenone exposure led to the appearance of donut/blob-shaped structures. In contrast, LM11A-31 completely prevented mitochondrial abnormalities (Figure 1F).

### 3.2 LM11A-31 hinders oxidative and nitrosative stress

Oxidative stress is one of the driving forces leading to dopaminergic neuronal death in PD. In this context, mitochondrial perturbances cause the premature transfer of electrons to O2 favoring the production of ROS and reactive nitrogen species (RNS) during oxidative phosphorylation. Given their strong tendency to rapidly react with various molecules, ROS and RNS can exert damage upon sub-cellular components (Singh et al. 2019). Thus, we evaluated oxidative damage to nucleic acids and lipids by using antibodies against 8-hydroxy-(deoxy)-guanosine (8-OH(d)G), 4-hydroxynonenal (4-HNE), and Nitro-tyrosine (N-Tyrosine) respectively. As expected, rotenone treatment led to increased oxidative and nitrosative damage that was normalized by p75NTR pharmacological modulation (Figure 2A-C).

**Figure 2.**
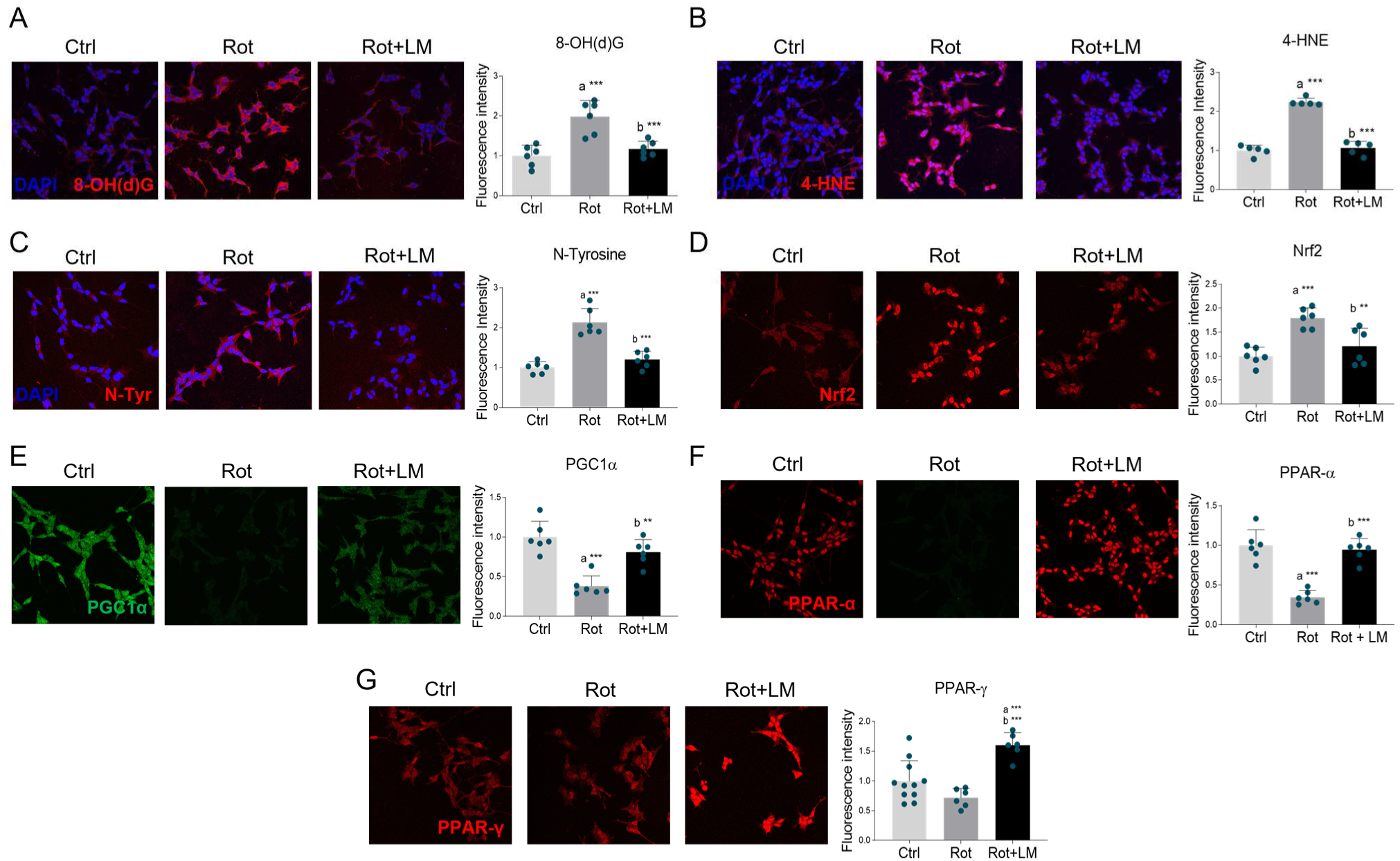
LM11A-31 effects on oxidative stress in rotenone-induced cell model of PD. Confocal microscopy representative images and fluorescent signal quantification of (A) 8-OH(d)G (red), (B) 4-HNE (red), (C) N-Tyrosine (red), (D) Nrf2 (red), (E) PGC1-α (green), (F) PPAR-α (red) and (G) PPAR-γ (red) in SH-SY5Y differentiated cell line treated with vehicle (Ctrl), 100 nM rotenone (Rot) and rotenone with 500 nM LM11A-31 (Rot+LM) for 24 h. DAPI (blue) was used to counterstain nuclei. N=5-11 independent experiments. Statistical analysis was performed by using one-way ANOVA followed by Tukey’s *post hoc* test. “a” indicates statistical significance vs Ctrl; “b” indicates statistical significance vs Rot group. ** p < 0.01, *** p < 0.001.

Redox homeostasis can be compromised when ROS production prevails over their catabolism. To cope with oxidative damage, cells employ antioxidant responses by inducing the transcription of enzymatic redox scavengers necessary for ROS detoxification. To understand the transcriptional mechanisms linking LM11A-31 to antioxidant activity, we examined the expression of the main master regulators in redox homeostasis. Nuclear factor erythroid 2-related factor 2 (Nrf2) is one of the best studied transcriptional factors, whose stabilization is increased by ROS accumulation (Fão et al. 2019). Accordingly, immunofluorescence analysis indicated that Nrf2 expression and nuclear localization were increased upon rotenone treatment, suggesting appropriate stress response. Interestingly, co-treatment with the p75NTR modulator normalized Nrf2 immunoreactivity, in line with the observation that LM11A-31 minimizes oxidative damage (Figure 2D). Redox balance also relies on other transcription factors entangled with antioxidant response, namely peroxisome proliferator-activated receptor (PPAR) α and γ and its co-activator peroxisome proliferator-activated receptor gamma coactivator 1-alpha (PGC-1α). Additionally, these proteins also contribute to mitochondrial and peroxisomal biogenesis, as well as organelle renewal (Scarpulla et al. 2011; Wójtowicz et al. 2020). Rotenone treatment markedly downregulated both PGC-1α and PPARα expression without significantly affecting PPARγ immunoreactivity, thus suggesting impaired antioxidant response. However, LM11A-31 boosted the expression of all these transcription factors (Figure 2E-G).

Aside from mitochondria, cells display other endogenous sources of ROS production, of which NADPH oxidase (NOX) is the most studied in PD pathogenesis (Pal et al., 2016). Even though no variations were observed in the expression of NOX2 and NOX4 catalytic subunits (figure 3A,B), rotenone exposure increased the immunopositivity associated with p22^PHOX^ and p47^PHOX^ regulatory subunits. Conversely, LM11A-31 significantly downregulated their expression (Figure C,D).

**Figure 3.**
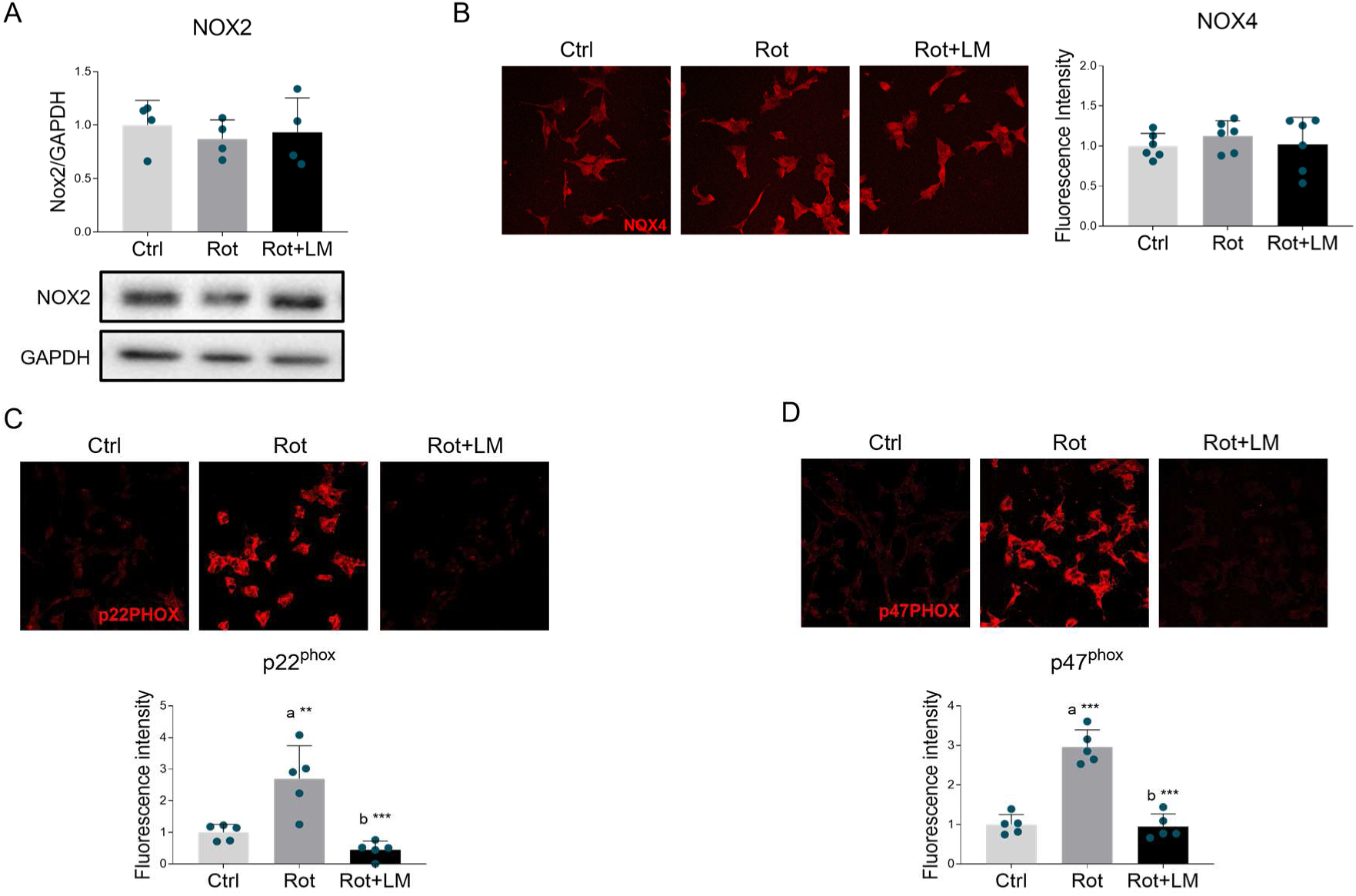
Effects of p75NTR modulation on NADPH oxidase complex in rotenone-treated SH-SY5Y cells. **(A)** Representative Western blot and densitometric analysis of NOX2 subunit in differentiated SH-SY5Y treated with DMSO (Ctrl), 100 nM rotenone (Rot) or rotenone with 500 nM LM11A-31 (Rot+LM) for 24h. GAPDH was chosen as loading control. N=4 biological replicates. (B-D) Representative immunofluorescence images and respective signal quantification of (B) NOX4, (C) p22phox and (D) p47phox in SH-SY5Y neuronal cells treated as in (A). N=5-6 independent experiments. Data are expressed as mean ± SD. Statistical analysis was assessed using the one-way ANOVA test, followed by Tukey’s *post hoc* test. Statistical significance is indicated as follows: ** p < 0.01; *** p < 0.001. “a” indicates statistical significance vs Ctrl group; “b” indicates statistical significance vs Rot group.

Taken together, these findings indicate that p75NTR pharmacological manipulation counteracts oxidative/nitrosative damage by reducing the expression of pro-oxidant enzymes. Furthermore, the increase in PPARs and PGC1α could contribute to enhance the antioxidant response and preserve mitochondrial integrity in rotenone-induced model of PD.

### 3.3 p75NTR modulation reverses rotenone-associated cholesterol dysmetabolism

Recent findings demonstrated that rotenone exposure induces profound alterations in cholesterol metabolism, downregulating the expression of NPC1 and subsequently favoring cholesterol build-up into lysosomes (Caria et al. 2023). In line with previous evidence, we observed free cholesterol accumulation in rotenone-treated cells, which was attenuated by LM11A-31 stimulation (Figure 4A). To evaluate whether the effects on free cholesterol could be dependent on intracellular cholesterol trafficking, we performed Western blot and immunofluorescence analysis, showing that NPC1 expression and localization to lysosomes were significantly reduced upon rotenone exposure; conversely, p75NTR modulation counteracted these alterations (Figure 4B,C). Cholesterol accumulation in rotenone-treated cells might also be attributed to the enhancement of lipoprotein uptake, as suggested by the significant increase of low-density lipoprotein receptor-related protein 1 (LRP1), which was normalized by LM11A-31 (Figure 4D). To further investigate the molecular mechanisms controlling cholesterol metabolism, we estimated the expression of the main enzymes involved in the biosynthetic pathway. The levels of the key and rate-limiting enzyme 3-hydroxy-3-methylglutaryl-CoA reductase (HMGCR) were reduced in rotenone-treated cells and restored upon p75NTR modulation (Figure 4E). Differently, no significant changes were observed in the expression of squalene synthase (SQS) and squalene epoxidase (SQE) (Figure 4F,G). Moreover, we performed immunofluorescence analysis on sterol regulatory element-binding protein 2 (SREBP-2), a master regulator of cholesterogenic genes. Interestingly, rotenone treatment significantly downregulated the expression of this transcription factor, which was unaffected by LM11A-31 (Figure 4H).

**Figure 4.**
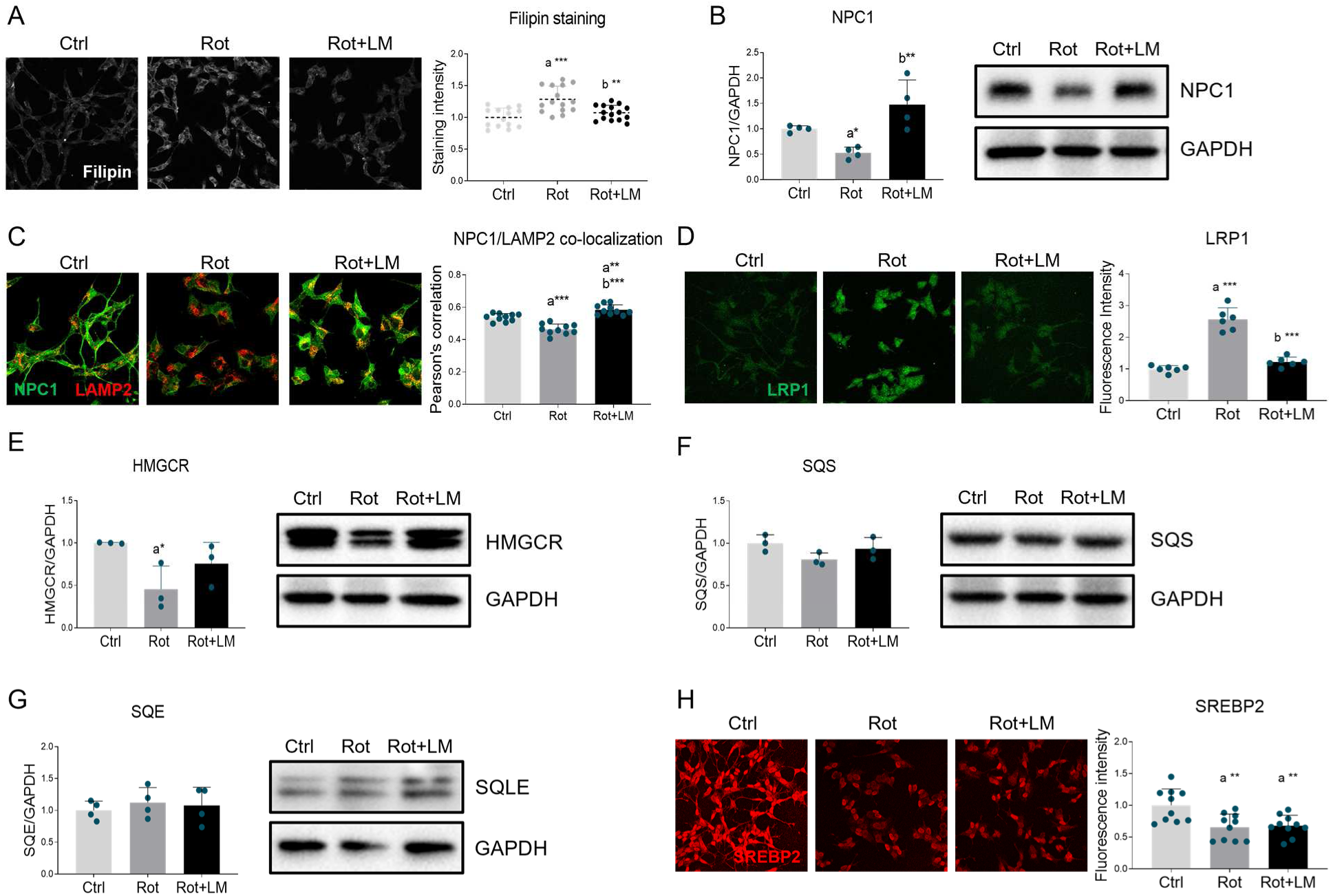
Impact of LM11A-31 on cholesterol metabolism in rotenone-treated cells. (A) Representative images (left panel) and quantification of filipin staining (right panel) performed on differentiated SH-SY5Y cells treated with vehicle (Ctrl), 100 nM rotenone (Rot) and rotenone with 500 nM LM11A-31 (Rot+LM) for 24 h. N=15 biological replicates. (B) SH-SY5Y treated as in A were subjected to Western blot and densitometric analysis for NPC1. GAPDH was used as housekeeping protein for loading control. N=4 biological replicates. (C) Co-immunofluorescence analysis performed on SH-SY5Y neuronal cells treated as previously reported, by using antibodies against NPC1 (green) and LAMP2 (red). Pearson’s correlation coefficient (r) was calculated to evaluate the protein co-localization. N=10 independent experiments. (D) Representative immunofluorescence images of LRP1 protein (green) in differentiated SH-SY5Y treated as previously described. N=6 biological replicates. (E-G) Representative immunoblot and densitometric analysis of HMGCR, SQS and SQE. GAPDH was used as internal loading control. N=3-4 independent experiments. (H) Immunofluorescence for SREBP2 performed of SH-SY5Y cells. N=10 biological replicates. Data are represented as means ± SD. Statistical analysis was performed by using one-way ANOVA followed by Tukey’s post hoc test. * p < 0.05, ** p < 0.01, *** p < 0.001. “a” indicates statistical significance vs Ctrl; “b” indicates statistical significance vs Rot group.

## Discussion

PD is the second most common neurodegenerative disorder after AD and it is mainly characterized by the loss of dopaminergic neurons in the *SNpc*. In addition to decreased dopamine levels in the *striatum*, several other pathological hallmarks have been identified, including protein aggregation, mitochondrial disfunctions and autophagy derangements (Liu et al. 2021; Ibarra-Gutiérrez et al. 2023). Along with these features, preclinical evidence indicated that p75NTR expression in dopaminergic neuronal cells is enhanced upon redox imbalance in PD, where it promotes α-syn aggregation and cell death (Chen et al. 2018). Given its ability to mediate both pro-survival and pro-apoptotic signaling cascades (Colardo et al. 2021), p75NTR has been widely considered as a valuable therapeutic target to counteract neurodegenerative conditions. In fact, several research groups have synthetized small molecules capable of mimicking mature neurotrophin binding to the receptor (i.e. LM11A-31), hence favoring cell viability over death (Massa et al., 2006). Currently, L-DOPA is the best therapeutic option for PD patients as it partially restores dopamine concentration in the *SNpc*. However, due to pharmacological tolerance over time and notorious side-effects such as L-DOPA-induced dyskinesia, new therapeutic avenues must be explored (Tran et al. 2018). Here, we display evidence that the pharmacological modulation of p75NTR by LM11A-31 prevents parkinsonian features in rotenone-treated SH-SY5Y cells. Immunofluorescence analysis first revealed p75NTR upregulation upon rotenone exposure, corroborating previous findings on receptor aberrations in PD (Chen et al. 2008). LM11A-31 exposure favored cell viability and reduced the number of apoptotic cells along with the rescue of neuromorphological aberrations and α-syn downregulation, when compared to rotenone-treated cells. Furthermore, LM11A-31 prevented mitochondrial fragmentation into donut/blob-like structures while concurrently contributing to reduce oxidative stress damage to macromolecules. These beneficial effects may be attributed to a boost in the expression of master regulators of the antioxidant response and mitochondrial homeostasis, namely PPARs and PGC1-α. Interestingly, Nguyen et al. demonstrated that 2-weeks treatment with LM11A-31 in stroked-mice elicited an increase in the available glutathione pool (Nguyen et al. 2022), probably indicating scavenging mechanism of the antioxidant response mediated by p75NTR modulation. Within cells, there are several sources of ROS production, such as damaged mitochondria, by-products of long-chain fatty acid β-oxidation and the pro-oxidant system such as NOX complexes. Keeny and colleagues highlighted that NOXs aberrations are critical events in the early stages of PD, as NOX2 activity is responsible for oxidative stress-dependent post-translational modification on α-syn resulting in its aggregation (Keeney et al. 2022). Our data corroborated this notion by showing increased expression of p22^PHOX^ and p47^PHOX^ regulatory subunits in rotenone-treated cells, which were normalized upon LM11A-31 treatment. It is therefore plausible to speculate that LM11A-31 effects on redox metabolism are not only based on the elevation of the antioxidant response, but also on diminishing ROS production by suppressing NOX activity, as recently evidenced in other pathological contexts (Varone et al., 2024).

Cholesterol is a critical component of biological membranes contributing to structural integrity and stability, supporting synapse formation and neurotransmitter release. In line with previous evidence (Caria et al. 2023), we reported intracellular cholesterol overload with reduced NPC1 expression and lysosomal localization, indicating derangements of cholesterol trafficking in PD. Additionally, HMGCR expression levels were found to be diminished in rotenone-treated cells, suggesting that reduced sterol biosynthesis may be a feedback response compensating the cholesterol intracellular overload. However, LM11A-31 normalized these alterations by restoring both NPC1 expression and lysosomal localization. Interestingly, it has previously been demonstrated that LM11A-31 exerts a regulatory effect on cholesterol metabolism. For instance, in astrocyte-like cells, LM11A-31 significantly induces HMGCR expression (Colardo et al., 2022). Even though further studies are required to better dissect the molecular mechanisms, it can be speculated that p75NTR modulation may rescue cholesterol redistribution in the cell, thus ameliorating lysosomal availability for protein aggregates and damaged-organelle degradation (Liao et al., 2007).

According to our findings, an elegant work has demonstrated that mitochondrial dysfunction, caused by complex I inhibition, elevates intracellular free cholesterol, which in turn suppresses SREBP-2 activation, leading to the downregulation of mevalonate pathway genes such as HMGCR. Besides altering transcriptional activity, the excess cholesterol induced by rotenone treatment also downregulates HMGCR expression by increasing its degradation (Wall et al., 2022). In addition, other reports have indicated that hypercholesterolemia, as well as cholesterol accumulation into lysosomes, alters physical properties of mitochondrial membranes, contributing to ROS leakage and subsequent spread of oxidative stress (Torres et al., 2017; Alrouji et al. 2023). Collectively, these lines of evidence suggest that cell dysfunction in PD may rely on a direct interplay between cholesterol homeostasis and redox balance, and that the cytoprotective effects promoted by LM11A-31 may be elicited by restoring these metabolic processes.

It is worth noting that LM11A-31 successfully entered phase IIa clinical trial for AD, showing adequate safety profile and bioavailability (Shanks et al. 2024). Additionally, LM11A-31 has been shown to restore Akt expression in the *striatum* while inhibiting c-Jun N-terminal kinases (JNK) pathways in R6/2 animal models, resulting in increased dendritic spine density and reduced inflammation (Simmons et al. 2016). These notions, along with the results shown in this work, suggest that p75NTR pharmacological modulation might be a valuable therapeutic approach not only in counteracting AD but also in other neurodegenerations such as PD and HD. However, other *in vitro* studies are certainly needed to better address the underlying molecular events and to identify the downstream effectors of the antioxidant response and pro-oxidant system in terms of both expression and activity. Furthermore, while cell culture models offer a simplified understanding of the pathology, future research is necessary to validate our findings through *in vivo* studies.

## Author contributions

Conceptualization: D.P. and M.S.; Data curation: D.P., M.S. and N.M.; Formal Analysis: D.P., M.S. and N.M; Funding acquisition: M.S.; Investigation: D.P., E.B.; F.C., G.S., M.C., M.V. and N.M.; Methodology: D.P., M.S. and N.M.; Project administration: D.P., M.S. and N.M.; Resources: M.S., S.M.; Software: M.S., S.M.; Supervision: M.S.; Validation: M.S., S.M.; Visualization: D.P., M.S.; Writing – original draft: D.P. and M.S.; Writing – review & editing: D.P., E.B.; F.C., G.S., M.C., M.S., M.V., N.M. and S.M.

